# Oligomerization-Mediated Phase-Separation in the Nucleoid-Associated Sensory Protein H-NS is Controlled by Ambient Cues

**DOI:** 10.1101/2024.08.10.607472

**Authors:** Bincy Lukose, Saloni Goyal, Athi N. Naganathan

## Abstract

H-NS, a nucleoid-associated protein (NAP) from enterobacteria, regulates gene expression by dynamically transducing environmental cues to conformational assembly and DNA binding. In this work, we show that H-NS from *Escherichia coli*, which can assemble into octameric and tetrameric oligomerization states, forms spontaneous micron-sized liquid-like condensates with DNA at sub-physiological concentrations *in vitro*. The heterotypic condensates are metastable at 298 K, partially solubilizing with time, while still retaining their liquid-like properties. The condensates display UCST-like phase behavior solubilizing at higher temperatures, but with a large decrease in droplet-assembly propensities at 310 K and also at higher ionic strength. Condensate formation can be tuned in a cyclic manner between 298 and 310 K with the extent of reversibility determined by the incubation time, highlighting strong hysteresis. An engineered phospho-mimetic variant (Y61E) of H-NS, which is dimeric and only weakly binds DNA, is unable to form condensates. The Y61E mutant solubilizes pre-formed H-NS condensates with DNA in a few minutes with nearly an order of magnitude speed-up in droplet dissolution at 310 K relative to 298 K, demonstrating rapid molecular transport between dilute and condensed phases. Our results establish that the oligomerization of H-NS is intrinsically tied not only to DNA binding but also its phase-separation tendencies, while showcasing the regulatable and programmable nature of heterotypic condensates formed by an archetypal NAP via multiple cues and their lifetimes.

## Introduction

Proteins of the H-NS family in bacteria play pivotal roles in transcription silencing involving pathogenic genes, environmentally regulated gene expression, genomic compaction, and xenogenic silencing.^1–7^ The native ensemble of H-NS from enterobacteria, and particularly from uropathogenic *E.coli*, is composed of an array of oligomeric species apart from folded monomeric and partially structured states.^8–15^ On changes in ambient conditions, be it changes in temperature, osmolarity, or pH, the extent of oligomerization is modulated which in turn controls not just the degree of DNA binding, but also the nature of assemblies that are formed. H-NS family members can also form linear lattices on dsDNA and even ‘bridged’ complexes wherein a linear array of oligomerized H-NS is sandwiched between two dsDNA molecules.^16–18^

The sheer diversity of regulatory processes governed by the small 137-residue, non-spherical, helical protein from uropathogenic *E.coli* (termed as H-NS from here on) has its origins in its architecture, with domains for dimerization, higher-order oligomerization and DNA-binding (Figure 1A, 1B). H-NS has a net charge of -2 despite harboring 48 charged residues (Figure 1A). The charged residues are also uniformly distributed with a κ-value, a sequence-based measure of charge-segregation,^19^ of just 0.2. The dimerization and oligomerization sites are both sequentially and structurally separate but with long-range allosteric connectivity.^15^ In addition, a 15-residue disordered linker separates the oligomerization domain from the DNA-binding domain (DBD). The surface charge distribution on the folded structure contributes to a positive potential on one face of the protein which is more evident when considering the tetrameric assembly (Figure 1C). We had shown earlier that the oligomerization of H-NS into tetramers and octamers is critical for efficient DNA binding.^15^ Specifically, a phosphomimetic mutation at position 61 (i.e. Y61E) disrupts the oligomeric interface stabilizing only the dimeric species in solution, which is unable to strongly bind DNA (Figure 1D).

**Figure 1.**
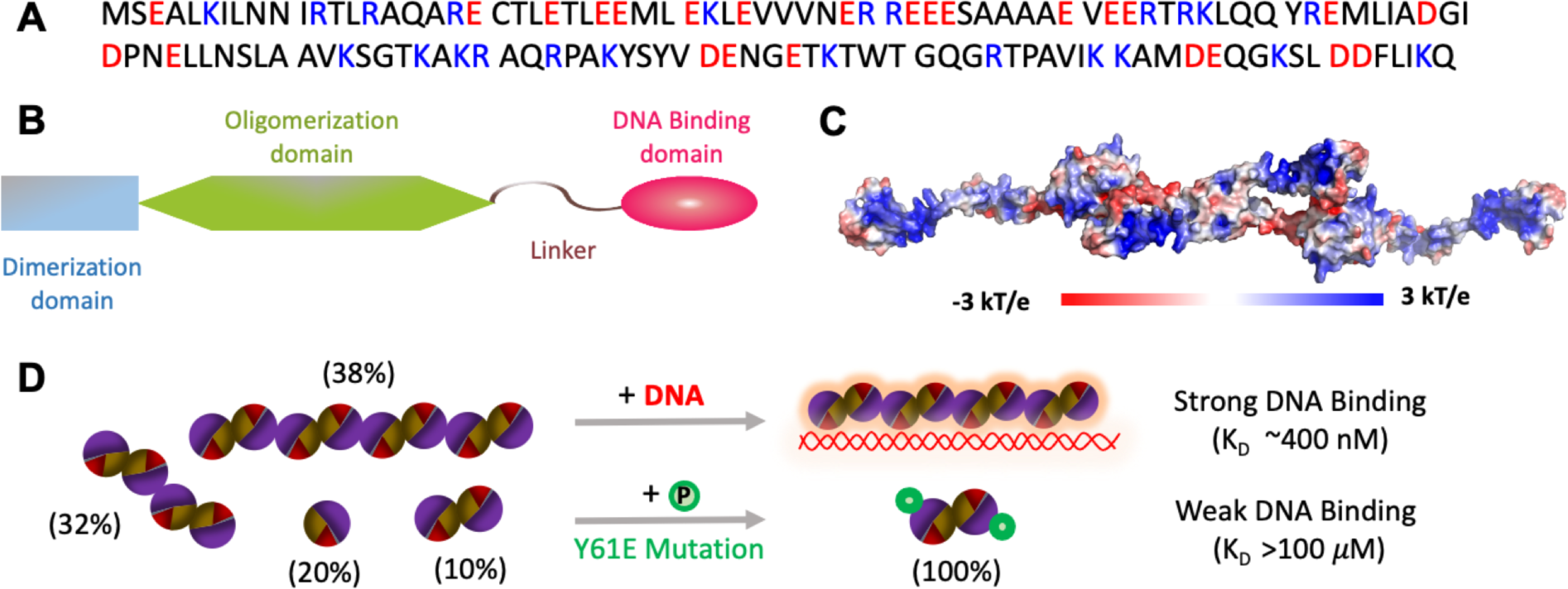
Structure of H-NS, oligomerization and DNA-binding. (A, B) Sequence of *E.coli* H-NS (panel A) and its domain organization (panel B). Positively and negatively charged residues are colored in blue and red, respectively, in panel A. (C) Electrostatic potential map^22^ of a tetrameric H-NS assembly model predicted from the AlphaFold2-ColabFold server.^23,24^ (D) The native ensemble of H-NS is populated by a collection of molecular species with different oligomerization status. Strong DNA binding is primarily via the tetrameric and octameric assembly modes (only the octameric binding mode is shown for clarity). The phospho-mimetic mutation (Y61E) eliminates higher-order oligomeric assemblies and the native ensemble is populated only by the dimer, which binds very weakly to DNA.

H-NS is therefore intrinsically multivalent in nature with the ability to form higher-order assemblies, with every protomer composed of both ordered and disordered regions. One could define the different binding domains and the associated positively charged patches as ‘stickers’ and disordered linker region as a ‘spacer’,^20,21^ a signature feature of many proteins that undergo phase separation.^25^ Given the structural complexity, multivalency, the function of H-NS, i.e. to condense bacterial DNA through non-specific weak interactions, and the fact that many bacterial proteins associated with DNA undergo phase separation,^26–29^ we hypothesize that H-NS is a prime candidate for forming phase-separated condensates with DNA. In fact, the moniker H-NS stands for histone-like nucleoid structuring, and is one member in the class of nucleoid-associated proteins (NAPs), apart from HU, IHF, Dps and FIS that constitute the bacterial nucleoid, which is a membraneless organelle housing the bacterial chromosome.^30–37^

In this work, we combine turbidity measurements, microscopy, and kinetic assays to show that H-NS forms condensates with DNA in a dose-dependent manner and at sub-physiological concentrations. The condensates are metastable displaying time-dependent solubilization while being intrinsically sensitive to both temperature and ionic strength, two factors that are known to strongly modulate bacterial gene expression through the disassembly of H-NS oligomers. Finally, through studies on the Y61E variant, we demonstrate that phase separation is inherently coupled to oligomerization, thus showcasing a potential regulatory mechanism to locally minimize the extent of condensate formation and hence liberate DNA for active transcription.

## Methods

### Purification of H-NS and Y61E

The proteins were purified as described before.^15^ All experiments reported here are carried out in 20 mM sodium phosphate buffer with sodium chloride salt (such that the effective ionic strength is 150 mM) at pH 8.0.

### Turbidity measurements

Turbidity measurements at 298 K were performed in a multimode plate reader (Agilent Synergy BioTek) in continuous mode to map the range of conditions under which H-NS undergoes phase separation. Different concentrations of H-NS ranging from 50 nM to 100 µM were dissolved in sodium phosphate buffer of pH 8 with or without 100 bp AT-rich DNA^14,15^ (400 nM) for a total volume of 150 *μ*l and loaded into a 96-well plate. The optical path length for this volume is estimated 3.94 mm. The turbidity of these solutions was monitored at 350 nm for 8 hours. Subsequent microscopy experiments were conducted with a fixed concentration of H-NS (100 µM) and a DNA concentration of 400 nM in 150 mM buffer of pH 8. Similar conditions were employed for the Y61E variant with 400 nM DNA in the same buffer. The resulting OD is reported without correction for optical path length. The experiments were carried out in duplicates and reported as mean ± s.d. (standard deviation).

To investigate the effect of temperature on condensates, turbidity was monitored at 350 nm as a function of temperature in a JASCO V-760 UV visible spectrophotometer equipped with a Peltier system in a 1 cm pathlength cuvette. The data was recorded from 5 – 95 ℃ with an increment of 1 ℃, and at a scan rate of 1 ℃/minute. Three scans were carried out in the temperature-dependent turbidity measurements, and the data are reported as mean ± s.d. All experiments, except for fluorescence microscopy, were carried with the unlabeled protein.

### Rhodamine labeling

For fluorescence microscopy experiments, the H-NS was labeled with NHS-Rhodamine (ThermoFisher Scientific, USA). Labeling was done at a dye-to-protein molar ratio of 5:1. Rhodamine was dissolved in DMSO and added to the protein in a conjugation buffer (0.1 M sodium phosphate, 0.15 M NaCl; pH 7.4). The protein-dye mixture was incubated at room temperature in a shaker with slow stirring for 2 hours. The unbound dye was removed by desalting in 150 mM buffer. The labeled protein concentration was calculated after diluting the sample as per the manufacturer’s protocol. After desalting, the labeled protein solution was stored at -20 ℃. For fluorescence microscopy experiments, 1:20 (v/v) of labeled versus unlabeled protein was used.

### Fluorescence microscopy

Microscopy-based experiments were carried out in an Olympus IX83 inverted fluorescence microscope. Glass slides and coverslips (Blue Star, India) were treated with piranha solution (3:1 v/v H_2_SO_4_:H_2_O_2_) to remove organic impurities. The treated slides were placed in a fume hood for 40 minutes and then thoroughly rinsed with Milli-Q water. Subsequently, the slides were cleaned with 100% ethanol and allowed to air-dry. The protein-DNA mixture was drop-casted on the acid-treated slide and covered with an acid-treated glass coverslip of 0.17 mm thickness. The droplets were visualized using a 100X oil immersion objective in either Differential Interference Contrast (DIC) or Fluorescence modes using an appropriate fluorescence channel. Droplet formation was monitored at different time points and different protein concentrations. The average diameter and number of the droplets were calculated from images by using in-house MATLAB codes. All images were captured with a 2048 x 2048 pixels resolution, except for the experiments with PEG8000 which employed 1024 x 1024 pixels resolution.

### Fluorescence recovery after photobleaching (FRAP)

The FRAP analysis of droplets was performed using a laser scanning Olympus Fluoview 3000 confocal microscope. Fluorescence intensity measurements were taken from five distinct regions of interest (ROIs) with consistent radii - the bleaching region (ROI-1), a reference region located on a neighboring droplet (ROI-2) to account for passive bleaching during excitation, and three regions outside the droplet in a dark area (ROI-3, -4, -5) to correct for background fluorescence intensity. ROI-1, underwent bleaching using a 561 nm laser channel, while emission intensities from all ROIs were monitored for ∼90 seconds until fluorescence emission from ROI-1 plateaued. The raw data was analyzed following the equation below^38^

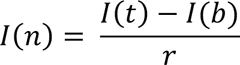

where *I*(*t*) is the fluorescence intensity at time *t*, *I*(*b*) is the background fluorescence intensity (average of ROI-3, -4, -5) and *r* is the ratio defined as *I*_*C*_/*I*_*C*0_. In the latter ratio, *I*_*C*0_ is the fluorescence intensity of the ROI-2 before photo-bleaching and *I*_*C*_ is the fluorescence intensity of the ROI-2 after photo-bleaching. The data were fit to single-exponential functions to extract the mean life-time and hence the recovery half-time (*t*_1/2_).^39^ FRAP experiments were carried out on three different droplets at every observation time point at [H-NS]:[DNA] of 50 *μ*M:400 nM, their profiles averaged and reported as mean ± s.d. (standard deviation).

### Stopped-flow experiments

Kinetics of condensate dissolution was monitored at 350 nm using a Chirascan SF.3 stopped-flow apparatus attached to the Chirascan qCD spectrometer (Applied Photophysics Ltd.) and a thermostated water bath. Y61E + DNA was taken in one 1 ml syringe while H-NS + DNA was loaded in another syringe and mixed rapidly (dead-time of ∼3 milliseconds). The resulting loss in OD due to droplet dissolution was followed as a function of time at 298 and 310 K in a 10 mm pathlength cell. The concentration of DNA was fixed at 400 nM in both the syringes so that the total [DNA] does not change on mixing. The final H-NS concentration was 100 *μ*M in the reaction mixture with different concentrations of Y61E including 50 *μ*M, 100 *μ*M and 200 *μ*M resulting in H-NS:Y61E molar ratios of 1:0.5, 1:1 and 1:2, respectively.

## Results and Discussion

### H-NS forms condensates with DNA

The prior DNA-binding experiments on H-NS were carried out at 40 nM of dye-labeled 100-bp dsDNA under a range of protein concentrations.^15^ Here, we ask if there are any signs of H-NS forming condensates at higher [DNA] which is one order of magnitude higher; 400 nM of duplex DNA effectively corresponds to 0.029 mg/ml, which is significantly lower than the nucleoid concentration of 7.1 mg/ml.^40^ Indeed, we observe droplet-like features in differential interference contrast (DIC) microscopy at 400 nM of 100-bp DNA and 100 *μ*M H-NS at 298 K (Figure 2A). More direct evidence is seen in fluorescence microscopy of NHS-rhodamine-labeled H-NS with DNA, which demonstrates the presence of phase-separated condensed droplets with an average diameter of 5 microns (Figure 2B). The droplets are highly dynamic, displaying multiple coalescence events within a timescale of a few seconds, indicating that they are liquid-like in nature (see Figure 2C for a representative example and Movie S1). FRAP (fluorescence recovery after photobleaching) experiments on the Rhodamine-labeled protein result in nearly 84% recovery of fluorescence on photobleaching, with a half-life (*t*_1/2_) of 9.4 seconds within the first 10 minutes (Figure 2D).

**Figure 2.**
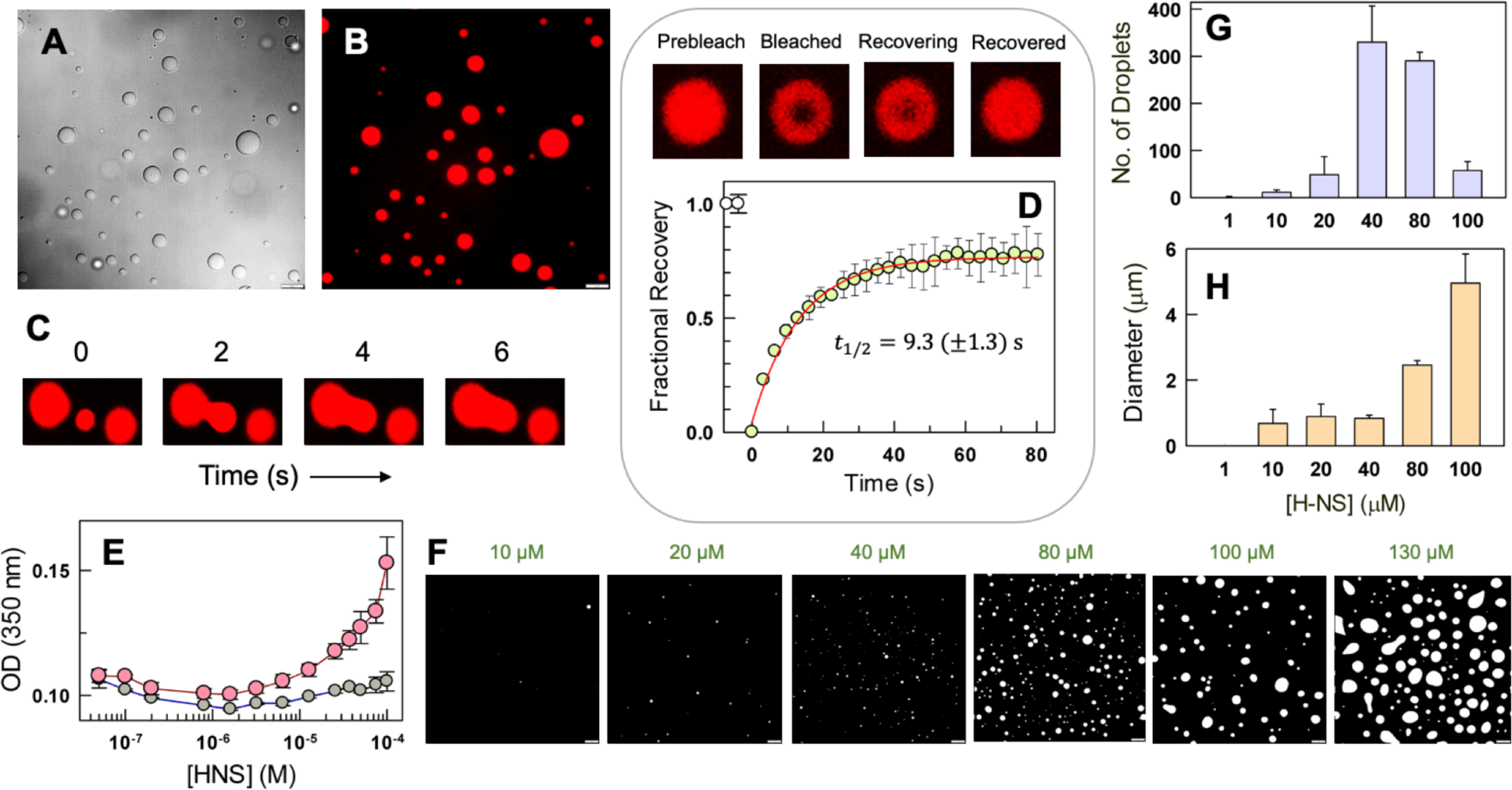
Associative phase transition of H-NS with DNA. (A, B) DIC and fluorescence microscopy images of NHS-Rhodamine labeled H-NS in the presence of 400 nM DNA. (C) A representative droplet fusion event observed via fluorescence microscopy. (D) Fluorescence microscopy images of time-dependent fluorescence recovery in condensates (circles) upon photobleaching. The recovery half-time extracted from a single-exponential fit to the data (red curve) is shown within the image. (E) Changes in turbidity as a function of increasing [H-NS] both in the presence (peach) and absence (gray) of 400 nM DNA. (F) Representative fluorescence images highlighting the H-NS concentration-dependence of phase separation. The droplets are shown in white for better contrast – this is followed in all fluorescence microscopy images from here on. (G, H) Number of droplets and droplet diameter as a function of [H-NS] in the presence of DNA. The scale bar at the bottom right of the microscopy images represents 10 microns.

In addition, we see a clear protein concentration dependence on droplet dimensions and numbers. The turbidity of H-NS mixture with DNA (at 400 nM) increases steeply beyond a [H-NS] of 10 *μ*M, while the reference protein solution displays little change (Figure 2E). Microscopy images illustrate that droplets can be observed at concentrations as low as 10-20 *μ*M of H-NS, much lower than the physiological concentration of ∼130 *μ*M.^41^ With increasing [H-NS], the number of droplets increase peaking at 40 *μ*M, beyond which it decreases due to coalescence of smaller droplets into larger assemblies (Figure 2F, 2G). This is observable in the plot of the diameter of the droplets that increase steeply from ∼1 *μ*m at 40 *μ*M to ∼5 *μ*m at 100 *μ*M (Figure 2H). Beyond 100 *μ*M, a strong wetting behavior is evident with irregular shaped droplets of sizes larger than 5 microns (not quantified due to the irregular shape, see the right-most panel in Figure 2F).

### H-NS condensates are metastable

The H-NS:DNA mixtures display a strong time-dependent decrease in turbidity in the first 20 minutes, following which it decreases slowly for nearly four hours. The turbidity then starts to increase again and with a similar time-dependence after 4 hours (peach in Figure 3A). The latter trend is likely a consequence of aggregation as solutions containing only H-NS display a similar time-dependence (gray in Figure 3A). Fluorescence microscopy images attest to the metastability, visually revealing a decrease in droplet density with time (Figure 3B, 3C). The metastability, defined here as the strong time-dependence, despite forming droplets spontaneously is quite surprising. Moreover, it is observable across two different approaches at a [H-NS] of 100 *μ*M which is nearly an order of magnitude higher than the saturation concentration (*C_sat_*) of <10 *μ*M expected from the concentration dependence studies (Figure 2F), and close to the physiological range of 130 *μ*M.^41^

**Figure 3.**
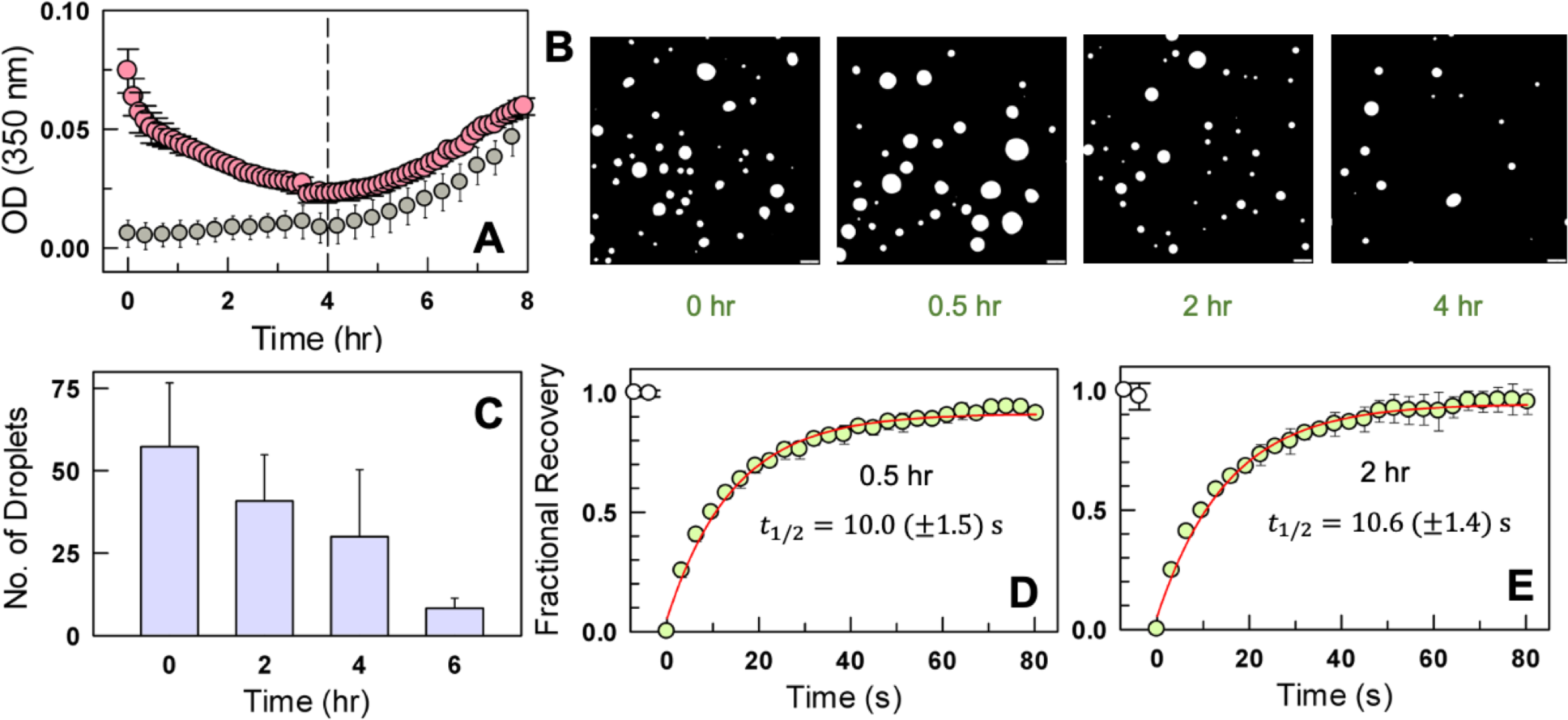
Metastability of H-NS condensates with DNA. (A) Turbidity changes as a function of time both in the presence (peach) and absence (gray) of 400 nM DNA at a fixed [H-NS] of 100 *μ*M (unlabeled H-NS). The vertical line indicates the timepoint till which the turbidity decreases (∼4 hrs). (B, C) Representative fluorescence microscopy images of NHS-rhodamine labeled H-NS:DNA mixtures at the indicated time-points highlighting the dissolution of droplets and the corresponding numbers (panel C) from a quantitative analysis of the images. Scale bars are 10 microns. (D, E) Fluorescence recovery time-profiles at 0.5 hours (panel D) and 2 hours (panel E) are shown in circles and the corresponding single-exponential fits in red. Note that the time-constants and the recovery amplitudes are similar to that acquired within a few minutes (Figure 2D), indicating that the droplets maintain their liquid-like character or a similar morphology until 2 hours despite solubilizing.

At 30 minutes and 2 hours, the FRAP recovery half-time is estimated to be only marginally longer at 10 and 10.6 seconds, respectively (compare Figure 3D, 3E with Figure 2D). The recovery extents (i.e. the amplitude) are marginally higher at longer times with nearly 90-95% recovery compared to ∼84% within the first 10 minutes. This is evidence that the droplets do maintain their liquid-like nature at longer observation times. These experiments also highlight that the condensate interface with the dilute phase is highly porous allowing a relatively rapid loss of protein (see the section on the Y61E mutant). It thus appears that the condensed phase is more kinetically accessible on mixing H-NS and DNA, despite not being the most stable phase under these conditions. This condensed phase undergoes compositional and potentially structural re-arrangements solubilizing with time while simultaneously seeding aggregation (the most stable phase) in the dilute phase at longer time-scales, as hypothesized in a recent review^42^ and observed experimentally in prion-like low complexity domains.^43^

### Condensates are sensitive to temperature variations

The metastability raises questions on the extent to which the droplet stability is dependent on temperature, a natural thermodynamic variable that strongly affects the assembly properties of free H-NS^15^ (i.e. in the absence of DNA). The turbidity of H-NS:DNA solution is high at low temperatures (<298 K), but with a strong temperature dependence while decreasing to baseline values at temperatures >315 K (blue in Figure 4A). The turbidity is only marginal at 310 K, signaling that the driving force for droplet formation is minimal at the optimal growth temperature of *E.coli*. Thus, the H-NS:DNA mixture exhibit properties of a UCST (upper critical solution temperature^21,44^) phase behavior with a cloud point temperature of 300 K (estimated from a first derivative of the OD versus temperature curve).

**Figure 4.**
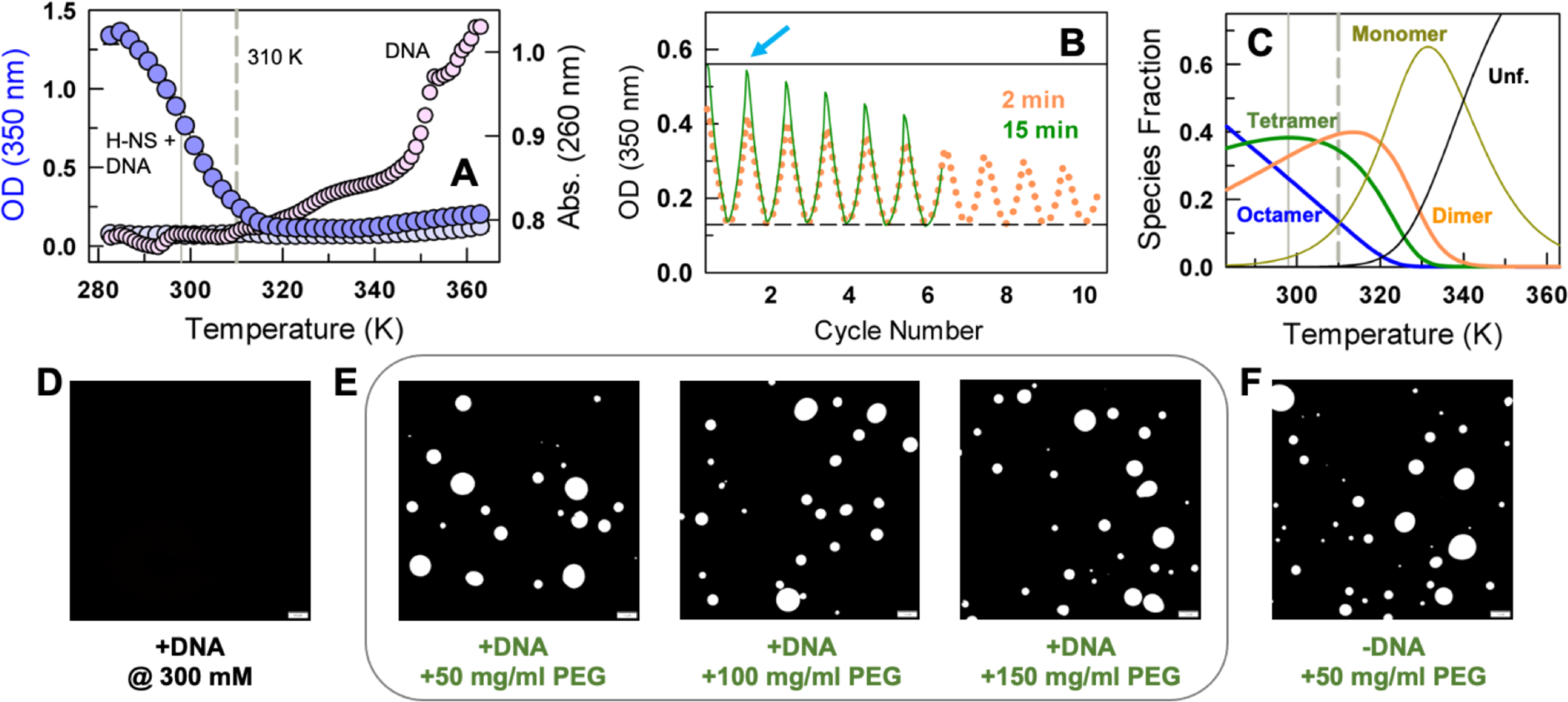
Effects of temperature, ionic strength and molecular crowding. (A) Changes in turbidity as a function of temperature at 150 mM (purple) and 300 mM ionic strength (gray) at 100 *μ*M H-NS and 400 nM DNA (left axis). The melting profile of DNA as monitored by absorbance at 260 nm (pink, right axis). The continuous and dashed vertical lines represent 298 and 310 K, respectively. (B) Changes in turbidity in a thermocycling experiment from 298 K (continuous horizontal line) to 310 K (dashed line) at different incubation times indicated and over multiple cycles at 150 mM ionic strength. The arrow represents the observation of full turbidity recovery in the first cycle and on 15 minutes incubation (green), while it never recovers fully at higher cycle numbers. (C) Predicted population of H-NS oligomeric species as a function of temperature from a thermodynamic model. The continuous and dashed vertical lines represent 298 and 310 K, respectively. (D) Representative microscopy image highlighting the absence of droplets at 300 mM ionic strength. (E) Droplets are observed even in crowding conditions at 100 *μ*M [H-NS] and 400 nM DNA. (F) H-NS phase separates into droplets upon moderate crowding and in the absence of DNA. Scale bars are 10 microns.

To test for reversibility of turbidity changes and potential hysteresis effects,^45^ we perform a thermocycling experiment wherein the H-NS:DNA mixture is heated at a scan rate of 1 °C/min from 298 K (high OD) to 310 K (low OD), and left to incubate for 2 minutes 310 K. Following this, the sample is cooled again at the same rate to 298 K and equilibrated again for 2 minutes. This procedure is repeated multiple times to explore the effects of repeated heating-cooling cycles on the condensate. A 2-minute equilibration time results in the final OD continually decreasing with repeated thermocycling, with only minimal OD recovery at the tenth cycle (orange in Figure 4B). On the other hand, a 15-minute equilibration in the thermocycling experiment leads to nearly 100% turbidity recovery at the end of the first cycle (arrow in Figure 4B) while it again decreases with cycle number, but with a weaker hysteresis behavior compared to the 2-minute equilibration experiment (green in Figure 4B). This experiment reveals that the dissolution of droplets can be quite quick within 2 minutes or less - similar to the timescale observed in protein-RNA-rich P granule dissolution *in vivo*^46^ - but making the intermolecular protein-DNA and protein-protein contacts necessary for effective droplet formation can be slow in this system. In addition, it is conceivable that an irreversible loss of H-NS to small aggregates or non-native conformations with every thermocycle contributes to the observed hysteresis.

The large temperature sensitivity at temperatures <310 K is not a consequence of structural changes in DNA, as the melting temperature of the 100 bp DNA we use is ∼340 K (pink in Figure 4A). In fact, through concentration-dependent sedimentation velocity experiments, heat capacity measurements, and a statistical thermodynamic model, we had shown earlier that the extent of H-NS oligomerization is temperature sensitive with the ensemble becoming monodisperse at higher temperatures.^15^ Employing the same thermodynamic model, we predict here that at a [H-NS] of 100 *μ*M, the native ensemble (in the absence of DNA) is dominated by octamers and tetramers at lower temperatures (Figure 4C). An increase in temperature leads to progressive disassembly of these oligomeric species with dimeric and monomeric H-NS populating at higher temperatures. Since the oligomerization of H-NS promotes stronger DNA binding, we hypothesize that this in turn promotes phase separation with DNA due to the build-up of the strong positive potential on one face of the H-NS assembly. At higher temperatures, the disassembly of oligomeric species leads to weaker interactions with DNA - as the dimeric variant of H-NS, Y61E, binds more than two orders of magnitude weaker to DNA - and hence reduces the propensity for phase separation under these conditions.

### Effects of ionic strength and crowding

Since H-NS is a non-specific DNA binding protein, the interactions are necessarily charge-mediated and sensitive to ionic strength.^47^ Increasing the solution ionic strength to 300 mM weakens the interactions with DNA by nearly two orders of magnitude (similar to the Y61E variant).^15^ The weakening of DNA binding at 300 mM ionic strength simultaneously eliminates phase separation in H-NS:DNA mixtures at the same concentration range as evidenced from both OD measurements (gray in Figure 4A) and fluorescence imaging (Figure 4D).

One final factor that is expected to affect the phase-separation behavior of H-NS is macromolecular crowding, which uniquely influences folding, stability and molecular diffusion in numerous systems.^48–51^ The experiments described so far were carried out in the absence of extrinsic crowding agents. On adding PEG-8000, a commonly used crowding agent, the phase separation properties of H-NS:DNA mixture are unchanged with condensates observed across the range of crowding conditions explored (Figure 4E). Surprisingly, H-NS by itself phase separates in the presence of 50 mg/ml of PEG8000 and without DNA. The driving force for phase-separation in such cases is expected to be different than the electrostatic complementarity driven complex coacervation observed in the presence of DNA,^52–54^ and hence we did not proceed further in our PEG-based studies.

### The dimeric Y61E H-NS variant solubilizes pre-formed H-NS condensates

Turbidity measurements and DIC/fluorescence microscopy demonstrate that the Y61E variant, a dimer and a weak DNA-binder, does not undergo phase separation with DNA at a range of protein concentrations, and incubation times (Figure 5A, 5B, S1). This result confirms that the driving force for phase separation in H-NS is oligomerization mediated strong non-specific DNA binding via electrostatic interactions.

**Figure 5.**
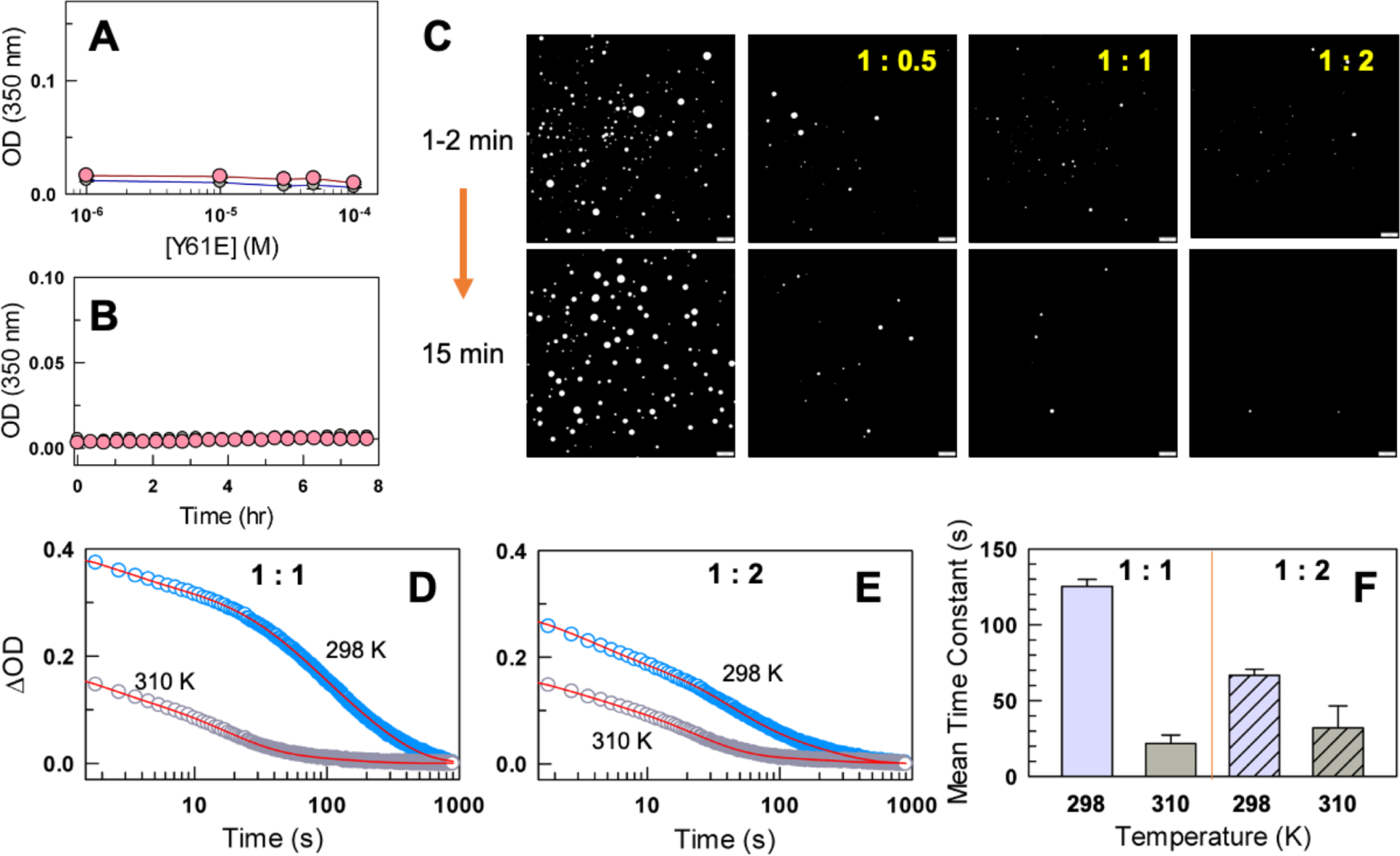
Porous nature of H-NS:DNA coacervates. (A, B) No changes in turbidity on incubating Y61E H-NS both in the presence (peach) and absence (gray) of 400 nM DNA (panel A). The turbidity does not vary even as a function of time (panel B, following color code of panel A). (C) Fluorescence microscopy images in the absence (left column) and at different indicated molar ratios of Y61E (right columns, H-NS:Y61E), right after mixing (within 1-2 minutes; top row) and 15-minutes after mixing (bottom row). The [H-NS] and [DNA] concentrations were fixed at 100 *μ*M and 400 nM, respectively. Scale bars are 10 microns. (D, E) Kinetic traces of changes in turbidity (ΔOD, circles) at different H-NS:Y61E molar ratios and temperatures, and the corresponding triple-exponential fits (red). (F) Amplitude-weighted time constants at 1:1 (colored bars) and 1:2 (colored and patterned bars) molar ratios.

To test if the Y61E variant can solubilize condensates, we add Y61E at different molar ratios – H-NS:Y61E of 1:0.5, 1:1, and 1:2 – to pre-formed H-NS:DNA condensates. Note that in these experiments, Y61E was also incubated with 400 nM DNA, so that the DNA concentration does not change on mixing. At H-NS:Y61E ratio of 1:0.5, a large reduction in droplet number and size can be observed (Figure 5C); the droplets persist even after 15 minutes, suggesting either that the timescale of solubilization is longer or that sub-stoichiometric concentrations of Y61E do not lead to complete dissolution. Fewer droplets persist at 1:1 ratio, while there is near-complete dissolution at 1:2 molar ratio. The solubilization of droplets on adding Y61E is a direct evidence to the liquid-like nature of the droplets and the highly porous nature of the condensate interface. The large dissolution even at sub-stoichiometric ratios of Y61E is because the dimerization (*D*) of H-NS is nearly two orders of magnitude stronger than oligomerization (*O*) as shown from earlier studies^15^: *K*_D_ is 500 nM while *K*_0_ is 30 *μ*M at 298 K. At 310 K, both weaken but with the dimer being more destabilized - *K*_D_ is 8 *μ*M while *K*_0_ is 45 *μ*M at 310 K – thus manifesting as a stronger dissolution effect.

To extract the associated timescales of droplet dissolution, we perform stopped-flow kinetics experiments, wherein a solution of H-NS:DNA is rapidly mixed (within a 3 milliseconds) with Y61E at different molar ratios (1:1 and 1:2) and two temperatures (298 and 310 K). The decrease in OD on droplet dissolution spans nearly three orders of magnitude (Figure 5D, 5E), thus necessitating at least two exponentials to describe the time-dependence. However, double-exponential fits were poorer, especially at the lowest time points (Figure S2). Tri-exponential fits provided very good fits with minimal fitting errors; each phase in the tri-exponential dependence is separated by an order of magnitude - a fast phase with a time constant of ∼1-3 seconds (across conditions), an intermediate phase with a time constant of 16-51 seconds, and a slow phase of 114-200 seconds (Table S1). The fast phase magnitude and its amplitude are nearly independent of the conditions, thus reporting on a fundamental process related to dissolution, the origins of which are unclear. The amplitudes and the time-constant of the intermediate phase increase and decrease with temperature, respectively, indicating that they describe a process related to molecular diffusivity. We do not attribute additional molecular meanings to the phases as the dissolution process is inherently complex coupled to the differential diffusivity of various oligomeric species in the dense phase and the dilute phase, in addition to the diffusivity of the different DNA-bound oligomeric states and their time dependent populations.

The mean time constant (*τ*), calculated as the weighted sum of the individual time constants and their amplitudes, however, speeds up by more than an order of magnitude at 310 K relative to 298 K at 1:1 molar ratio, while there is only a marginal speed-up of dissolution with temperature at 1:2 molar ratio (Figure 5F). If the faster dissolution at 310 K and at 1:1 molar ratio were to purely depend on changes in viscosity (which in turn affects the diffusion constants) between the two temperatures, then the *τ* at 310 K should be 97 seconds and not 21.8 seconds. Similarly, the correction for viscosity at 310 K (1:2 molar ratio) leads to a *τ* of 51.9 seconds, compared to the 32 seconds observed. This indicates that the faster dissolution at 310 K has additional contributions potentially from the temperature dependence of oligomeric species populations.

## Conclusions

There has been a growing interest in understanding the organization of bacterial nucleoid as it exists as a phase-separated blob within the cytoplasm harboring the bacterial chromosome, NAPs, RNA and RNA polymerase. H-NS is one of the many NAPs that bind and organize bacterial nucleoid. In this work, we show strong *in vitro* evidence that H-NS can condense DNA into phase-separated liquid-like droplets spontaneously and at sub-physiological protein and DNA concentrations as summarized in Figure 6. This observation adds on to a large body of work where electrostatic complementarity between the two macromolecules, protein and nucleic acids, is sufficient to drive an associative phase separation but under appropriate concentration regimes.^55,56,65–68,57–64^ The metastability, the molecular origins of which require further exploration, hints that H-NS by itself would not be sufficient to maintain the phase-separated behavior of the bacterial nucleoid. It is interesting to note that two other NAPs, HU and Dps both together and in isolation, are able to condense DNA^29^ in a manner similar to our observations. Given the large compaction of the bacterial chromosome into the nucleoid,^32,33^ it is very likely that many NAPs work together to maintain its liquid-like architecture. In such a scenario, the difference in binding affinity to DNA would effectively determine the different concentration regimes at which they bind and condense DNA.

**Figure 6.**
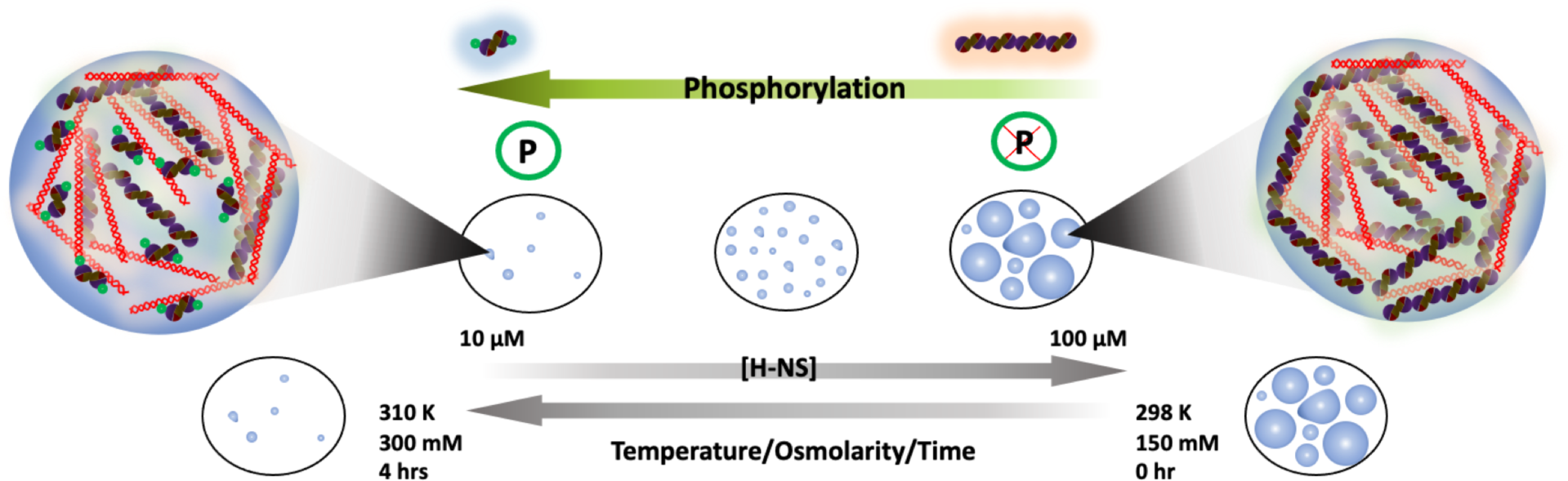
The native ensemble of H-NS is polydisperse with a large population of tetrameric and octameric assemblies, which in the presence of DNA phase separate into condensates with liquid-like character. The phosphomimetic variant, on the other hand, is unable to form condensates and is also able to solubilize pre-formed H-NS:DNA condensates. The droplets are highly metastable, being sensitive to time, small changes in temperature and osmolarity.

Proteomic studies on bacterial proteins have unearthed numerous post-translational modifications in bacterial NAPs, with H-NS alone undergoing PTMs at 29 different sites.^69^ The inability of the dimeric variant, Y61E, to condense DNA has important implications. Interestingly, a similar dimeric variant harboring two mutations Y61D/M64D has been shown to be less efficient in forming bridged complexes compared to the H-NS wild-type.^16^ Taken together with the results of the current work, we conclude that the condensation and phase-separation behavior of H-NS:DNA mixtures is controlled primarily by the oligomerization of H-NS. A direct test to this expectation comes from studies on the Y61E, which binds weakly to DNA as it exclusively populates only dimers in solution,^15^ and is unable to form condensates with DNA.

An open question in the field of biomolecular condensates is the nature of their regulation, i.e. can the condensates be solubilized to disassemble and release the constituents for subsequent functional requirements? With H-NS:DNA mixtures, we show that an increase in temperature reduces the extent of phase separation in the physiological range with only minimal turbidity observed at 310 K, highlighting that small modulations in thermal energy can have substantial effects on condensate formation propensities. This trait is again related to the sensitivity of H-NS oligomerization to temperature, as only dimers and tetramers (to a lower extent) are primarily populated at temperatures beyond 310 K. As the bacterial nucleoid has multiple proteins maintaining the liquid-like nature of the nucleoid,^35^ it is possible that the density of the nucleoid is modulated locally on increase in temperature due to H-NS disassembly (Figure 4C) and proportional loss of DNA-bridging. This would in turn relieve the transcription repression mediated by H-NS at 310 K by providing RNA polymerase the access to otherwise occluded promoter sites.

Finally, the ability of Y61E variant to rapidly solubilize pre-formed H-NS:DNA condensates brings to focus the porous nature of the droplet interface, which enables free exchange of macromolecules between the dilute and condensed phases. In the absence of cues related to temperature or osmolarity, an excess phosphorylated protein (mimicked in this work by the Y61E variant) can alone substantially solubilize the condensates within a few minutes. Each of the PTMs identified earlier^69^ could therefore modulate dimerization, oligomerization or DNA-binding equilibria distinctly regulating different facets of H-NS assembly and hence by extension phase separation with DNA. Our results thus put forth heterotypic coacervation of H-NS with DNA as one more layer of control to modulate the expression of genes in bacteria.

## Abbreviations

H-NS: histone-like nucleoid condensing
LLPS: liquid-liquid phase separation
DIC: differential interference contrast
DNA: deoxy ribonucleic acid
FRAP: fluorescence recovery after photobleaching

## Acknowledgements

The authors are grateful for the support from the Department of Biotechnology (DBT, India) for the grant BT/PR41973/BRB/10/1967/2021 to A. N. N. The authors acknowledge the FIST facility (SR/FST/LS-II/2020/552(C)) sponsored by the Department of Science and Technology (DST, India) and the ICSR - Common Instruments Facility (CIF) at the Department of Biotechnology, IIT Madras (Chennai, India) for the instrumentation. The authors thank Anup Kumar Mani for help with microscopy. B.L. thanks the funding from the Women Leading IITM initiative.

## REFERENCES

(1) Nieto, J. M.; Madrid, C.; Prenafeta, A.; Miquelay, E.; Balsalobre, C.; Carrascal, M.; Juarez, A. Expression of the Hemolysin Operon in Escherichia Coli Is Modulated by a Nucleoid-Protein Complex That Includes the Proteins Hha and H-NS. Mol. Gen. Genet. 2000, 263 (2), 349–358.

(2) Madrid, C.; Nieto, J. M.; Paytubi, S.; Falconi, M.; Gualerzi, C. O.; Juarez, A. Temperature- and H-NS-Dependent Regulation of a Plasmid-Encoded Virulence Operon Expressing Escherichia Coli Hemolysin. J. Bacteriol. 2002, 184 (18), 5058–5066.

(3) Banos, R. C.; Pons, J. I.; Madrid, C.; Juarez, A. A Global Modulatory Role for the Yersinia Enterocolitica H-NS Protein. Microbiology 2008, 154 (Pt 5), 1281–1289. 10.1099/mic.0.2007/015610-0.

(4) Grainger, D. C. Structure and Function of Bacterial H-NS Protein. Biochem. Soc. Transac. 2016, 44 (6), 1561–1569. 10.1042/BST20160190.

(5) Qin, L.; Erkelens, A. M.; Ben Bdira, F.; Dame, R. T. The Architects of Bacterial DNA Bridges: A Structurally and Functionally Conserved Family of Proteins. Open Biol. 2019, 9 (12), 190223. 10.1098/rsob.190223.

(6) Dame, R. T.; Rashid, F.-Z. M.; Grainger, D. C. Chromosome Organization in Bacteria: Mechanistic Insights into Genome Structure and Function. Nat. Rev. Genet. 2020, 21 (4), 227–242. 10.1038/s41576-019-0185-4.

(7) Rashid, F.-Z. M.; Dame, R. T. Three-Dimensional Chromosome Re-Modelling: The Integral Mechanism of Transcription Regulation in Bacteria. Mol. Microbiol. 2023, 120 (1), 60–70. 10.1111/mmi.15062.

(8) Renzoni, D.; Esposito, D.; Pfuhl, M.; Hinton, J. C.; Higgins, C. F.; Driscoll, P. C.; Ladbury, J. E. Structural Characterization of the N-Terminal Oligomerization Domain of the Bacterial Chromatin-Structuring Protein, H-NS. J. Mol. Biol. 2001, 306 (5), 1127–1137. 10.1006/jmbi.2001.4471.

(9) Esposito, D.; Petrovic, A.; Harris, R.; Ono, S.; Eccleston, J. F.; Mbabaali, A.; Haq, I.; Higgins, C. F.; Hinton, J. C.; Driscoll, P. C.; Ladbury, J. E. H-NS Oligomerization Domain Structure Reveals the Mechanism for High Order Self-Association of the Intact Protein. J. Mol. Biol. 2002, 324 (4), 841–850.

(10) Leonard, P. G.; Ono, S.; Gor, J.; Perkins, S. J.; Ladbury, J. E. Investigation of the Self-Association and Hetero-Association Interactions of H-NS and StpA from Enterobacteria. Mol. Microbiol. 2009, 73 (2), 165–179. 10.1111/j.1365-2958.2009.06754.x.

(11) Arold, S. T.; Leonard, P. G.; Parkinson, G. N.; Ladbury, J. E. H-NS Forms a Superhelical Protein Scaffold for DNA Condensation. Proc. Natl. Acad. Sci. U.S.A. 2010, 107 (36), 15728–15732. 10.1073/pnas.1006966107.

(12) Cordeiro, T. N.; Garcia, J.; Bernado, P.; Millet, O.; Pons, M. A Three-Protein Charge Zipper Stabilizes a Complex Modulating Bacterial Gene Silencing. J. Biol. Chem. 2015, 290, 21200–21212.

(13) Narayan, A.; Bhattacharjee, K.; Naganathan, A. N. Thermally versus Chemically Denatured Protein States. Biochemistry 2019, 58 (21), 2519–2523. 10.1021/acs.biochem.9b00089.

(14) Shahul Hameed, U. F.; Liao, C.; Radhakrishnan, A. K.; Huser, F.; Aljedani, S. S.; Zhao, X.; Momin, A. A.; Melo, F. A.; Guo, X.; Brooks, C.; Li, Y.; Cui, X.; Gao, X.; Ladbury, J. E.; Jaremko, Ł.; Jaremko, M.; Li, J.; Arold, S. T. H-NS Uses an Autoinhibitory Conformational Switch for Environment-Controlled Gene Silencing. Nuc. Acids Res. 2019, 47 (5), 2666–2680. 10.1093/nar/gky1299.

(15) Lukose, B.; Maruno, T.; Faidh, M. A.; Uchiyama, S.; Naganathan, A. N. Molecular and Thermodynamic Determinants of Self-Assembly and Hetero-Oligomerization in the Enterobacterial Thermo-Osmo-Regulatory Protein H-NS. Nuc. Acids Res. 2024, 52 (5), 2157–2173. 10.1093/nar/gkae090.

(16) van der Valk, R. A.; Vreede, J.; Qin, L.; Moolenaar, G. F.; Hofmann, A.; Goosen, N.; Dame, R. T. Mechanism of Environmentally Driven Conformational Changes That Modulate H-NS DNA-Bridging Activity. Elife 2017, 6. 10.7554/eLife.27369.

(17) Qin, L.; Bdira, F. B.; Sterckx, Y. G. J.; Volkov, A. N.; Vreede, J.; Giachin, G.; van Schaik, P.; Ubbink, M.; Dame, R. T. Structural Basis for Osmotic Regulation of the DNA Binding Properties of H-NS Proteins. Nuc. Acids Res. 2020, 48 (4), 2156–2172. 10.1093/nar/gkz1226.

(18) Shen, B. A.; Hustmyer, C. M.; Roston, D.; Wolfe, M. B.; Landick, R. Bacterial H-NS Contacts DNA at the Same Irregularly Spaced Sites in Both Bridged and Hemi-Sequestered Linear Filaments. iScience 2022, 25 (6), 104429. 10.1016/j.isci.2022.104429.

(19) Das, R. K.; Pappu, R. V. Conformations of Intrinsically Disordered Proteins Are Influenced by Linear Sequence Distributions of Oppositely Charged Residues. Proc. Natl. Acad. Sci. U.S.A. 2013, 110 (33), 13392–13397. 10.1073/pnas.1304749110.

(20) Cohan, M. C.; Pappu, R. V. Making the Case for Disordered Proteins and Biomolecular Condensates in Bacteria. Trends Biochem. Sci. 2020, 45 (8), 668–680. 10.1016/j.tibs.2020.04.011.

(21) Pappu, R. V; Cohen, S. R.; Dar, F.; Farag, M.; Kar, M. Phase Transitions of Associative Biomacromolecules. Chem.Rev. 2023, 123 (14), 8945–8987. 10.1021/acs.chemrev.2c00814.

(22) Dolinsky, T. J.; Nielsen, J. E.; McCammon, J. A.; Baker, N. A. PDB2PQR: An Automated Pipeline for the Setup of Poisson-Boltzmann Electrostatics Calculations. Nucleic Acids Res. 2004, 32 (Web Server issue), W665-7. 10.1093/nar/gkh381.

(23) Jumper, J.; Evans, R.; Pritzel, A.; Green, T.; Figurnov, M.; Ronneberger, O.; Tunyasuvunakool, K.; Bates, R.; Žídek, A.; Potapenko, A.; Bridgland, A.; Meyer, C.; Kohl, S. A. A.; Ballard, A. J.; Cowie, A.; Romera-Paredes, B.; Nikolov, S.; Jain, R.; Adler, J.; Back, T.; Petersen, S.; Reiman, D.; Clancy, E.; Zielinski, M.; Steinegger, M.; Pacholska, M.; Berghammer, T.; Bodenstein, S.; Silver, D.; Vinyals, O.; Senior, A. W.; Kavukcuoglu, K.; Kohli, P.; Hassabis, D. Highly Accurate Protein Structure Prediction with AlphaFold. Nature 2021, 596, 583–589. 10.1038/s41586-021-03819-2.

(24) Mirdita, M.; Schütze, K.; Moriwaki, Y.; Heo, L.; Ovchinnikov, S.; Steinegger, M. ColabFold: Making Protein Folding Accessible to All. Nat. Methods 2022, 19 (6), 679–682. 10.1038/s41592-022-01488-1.

(25) Brangwynne, C. P.; Eckmann, C. R.; Courson, D. S.; Rybarska, A.; Hoege, C.; Gharakhani, J.; Jülicher, F.; Hyman, A. A. Germline P Granules Are Liquid Droplets That Localize by Controlled Dissolution/Condensation. Science *(80-. ).* 2009, 324 (5935), 1729–1732. 10.1126/science.1172046.

(26) Ladouceur, A.-M.; Parmar, B. S.; Biedzinski, S.; Wall, J.; Tope, S. G.; Cohn, D.; Kim, A.; Soubry, N.; Reyes-Lamothe, R.; Weber, S. C. Clusters of Bacterial RNA Polymerase Are Biomolecular Condensates That Assemble through Liquid-Liquid Phase Separation. Proc Natl. Acad. Sci. U. S. A. 2020, 117 (31), 18540–18549. 10.1073/pnas.2005019117.

(27) Harami, G. M.; Kovács, Z. J.; Pancsa, R.; Pálinkás, J.; Baráth, V.; Tárnok, K.; Málnási-Csizmadia, A.; Kovács, M. Phase Separation by SsDNA Binding Protein Controlled via Protein-Protein and Protein-DNA Interactions. Proc Natl. Acad. Sci. U. S. A. 2020, 117 (42), 26206–26217. 10.1073/pnas.2000761117.

(28) Kozlov, A. G.; Cheng, X.; Zhang, H.; Shinn, M. K.; Weiland, E.; Nguyen, B.; Shkel, I. A.; Zytkiewicz, E.; Finkelstein, I. J.; Record, M. T. J.; Lohman, T. M. How Glutamate Promotes Liquid-Liquid Phase Separation and DNA Binding Cooperativity of E. Coli SSB Protein. J.Mol.Biol. 2022, 434 (9), 167562. 10.1016/j.jmb.2022.167562.

(29) Gupta, A.; Joshi, A.; Arora, K.; Mukhopadhyay, S.; Guptasarma, P. The Bacterial Nucleoid-Associated Proteins, HU and Dps, Condense DNA into Context-Dependent Biphasic or Multiphasic Complex Coacervates. J.Biol.Chem. 2023, 299 (5), 104637. 10.1016/j.jbc.2023.104637.

(30) Kleppe, K.; Ovrebö, S.; Lossius, I. The Bacterial Nucleoid. J.Gen.Microbiol. 1979, 112 (1), 1–13. 10.1099/00221287-112-1-1.

(31) Robinow, C.; Kellenberger, E. The Bacterial Nucleoid Revisited. Microbiol.Rev. 1994, 58 (2), 211–232. 10.1128/mr.58.2.211-232.1994.

(32) Fisher, J. K.; Bourniquel, A.; Witz, G.; Weiner, B.; Prentiss, M.; Kleckner, N. Four-Dimensional Imaging of E. Coli Nucleoid Organization and Dynamics in Living Cells. Cell 2013, 153 (4), 882–895. 10.1016/j.cell.2013.04.006.

(33) Le, T. B. K.; Imakaev, M. V; Mirny, L. A.; Laub, M. T. High-Resolution Mapping of the Spatial Organization of a Bacterial Chromosome. Science *(80-. ).* 2013, 342 (6159), 731–734. 10.1126/science.1242059.

(34) Wiggins, P. A.; Cheveralls, K. C.; Martin, J. S.; Lintner, R.; Kondev, J. Strong Intranucleoid Interactions Organize the Escherichia Coli Chromosome into a Nucleoid Filament. Proc. Natl. Acad. Sci. U. S. A. 2010, 107 (11), 4991–4995. 10.1073/pnas.0912062107.

(35) Monterroso, B.; Margolin, W.; Boersma, A. J.; Rivas, G.; Poolman, B.; Zorrilla, S. Macromolecular Crowding, Phase Separation, and Homeostasis in the Orchestration of Bacterial Cellular Functions. Chem.Rev. 2024, 124 (4), 1899–1949. 10.1021/acs.chemrev.3c00622.

(36) Walker, A. M.; Abbondanzieri, E. A.; Meyer, A. S. Live to Fight Another Day: The Bacterial Nucleoid under Stress. Mol.Microbiol. 2024. 10.1111/mmi.15272.

(37) Hustmyer, C. M.; Landick, R. Bacterial Chromatin Proteins, Transcription, and DNA Topology: Inseparable Partners in the Control of Gene Expression. Mol.Microbiol. 2024. 10.1111/mmi.15283.

(38) Ray, S.; Singh, N.; Kumar, R.; Patel, K.; Pandey, S.; Datta, D.; Mahato, J.; Panigrahi, R.; Navalkar, A.; Mehra, S.; Gadhe, L.; Chatterjee, D.; Sawner, A. S.; Maiti, S.; Bhatia, S.; Gerez, J. A.; Chowdhury, A.; Kumar, A.; Padinhateeri, R.; Riek, R.; Krishnamoorthy, G.; Maji, S. K. α-Synuclein Aggregation Nucleates through Liquid-Liquid Phase Separation. Nat.Chem. 2020, 12 (8), 705–716. 10.1038/s41557-020-0465-9.

(39) Taylor, N. O.; Wei, M.-T.; Stone, H. A.; Brangwynne, C. P. Quantifying Dynamics in Phase-Separated Condensates Using Fluorescence Recovery after Photobleaching. Biophys.J. 2019, 117 (7), 1285–1300. 10.1016/j.bpj.2019.08.030.

(40) Xiang, Y.; Surovtsev, I. V; Chang, Y.; Govers, S. K.; Parry, B. R.; Liu, J.; Jacobs-Wagner, C. Interconnecting Solvent Quality, Transcription, and Chromosome Folding in Escherichia Coli. Cell 2021, 184 (14), 3626–3642.e14. 10.1016/j.cell.2021.05.037.

(41) Ali Azam, T.; Iwata, A.; Nishimura, A.; Ueda, S.; Ishihama, A. Growth Phase-Dependent Variation in Protein Composition of the Escherichia Coli Nucleoid. J. Bacteriol. 1999, 181 (20), 6361–6370. 10.1128/JB.181.20.6361-6370.1999.

(42) Zhang, Z.; Huang, G.; Song, Z.; Gatch, A. J.; Ding, F. Amyloid Aggregation and Liquid-Liquid Phase Separation from the Perspective of Phase Transitions. J.Phys.Chem. B 2023, 127 (28), 6241–6250. 10.1021/acs.jpcb.3c01426.

(43) Das, T.; Zaidi, F.; Farag, M.; Ruff, K. M.; Messing, J.; Taylor, J. P.; Pappu, R. V; Mittag, T. Metastable Condensates Suppress Conversion to Amyloid Fibrils. bioRxiv. United States March 2024. 10.1101/2024.02.28.582569.

(44) Shapiro, D. M.; Ney, M.; Eghtesadi, S. A.; Chilkoti, A. Protein Phase Separation Arising from Intrinsic Disorder: First-Principles to Bespoke Applications. J.Phys.Chem. B 2021, 125 (25), 6740–6759. 10.1021/acs.jpcb.1c01146.

(45) Garcia Quiroz, F.; Li, N. K.; Roberts, S.; Weber, P.; Dzuricky, M.; Weitzhandler, I.; Yingling, Y. G.; Chilkoti, A. Intrinsically Disordered Proteins Access a Range of Hysteretic Phase Separation Behaviors. Sci. Adv. 2019, 5 (10), eaax5177. 10.1126/sciadv.aax5177.

(46) Fritsch, A. W.; Diaz-Delgadillo, A. F.; Adame-Arana, O.; Hoege, C.; Mittasch, M.; Kreysing, M.; Leaver, M.; Hyman, A. A.; Jülicher, F.; Weber, C. A. Local Thermodynamics Govern Formation and Dissolution of Caenorhabditis Elegans P Granule Condensates. Proc. Natl. Acad. Sci. U. S. A. 2021, 118 (37). 10.1073/pnas.2102772118.

(47) Record Jr., M. T.; Courtenay, E. S.; Cayley, D. S.; Guttman, H. J. Responses of E. Coli to Osmotic Stress: Large Changes in Amounts of Cytoplasmic Solutes and Water. Trends Biochem. Sci. 1998, 23 (4), 143–148.

(48) Fulton, A. B. How Crowded Is the Cytoplasm? Cell 1982, 30 (2), 345–347. 10.1016/0092-8674(82)90231-8.

(49) Zimmerman, S. B.; Minton, A. P. Macromolecular Crowding: Biochemical, Biophysical, and Physiological Consequences. Ann. Rev. Biophys. Biomol. Struct. 1993, 22, 27–65. 10.1146/annurev.bb.22.060193.000331.

(50) Sarkar, M.; Li, C.; Pielak, G. J. Soft Interactions and Crowding. Biophys. Rev. 2013, 5 (2), 187–194. 10.1007/s12551-013-0104-4.

(51) Wennerström, H.; Vallina Estrada, E.; Danielsson, J.; Oliveberg, M. Colloidal Stability of the Living Cell. Proc. Natl. Acad. Sci. U. S. A. 2020, 117 (19), 10113–10121. 10.1073/pnas.1914599117.

(52) Ghosh, A.; Mazarakos, K.; Zhou, H.-X. Three Archetypical Classes of Macromolecular Regulators of Protein Liquid-Liquid Phase Separation. Proc. Natl. Acad. Sci. U. S. A. 2019, 116 (39), 19474–19483. 10.1073/pnas.1907849116.

(53) Park, S.; Barnes, R.; Lin, Y.; Jeon, B.-J.; Najafi, S.; Delaney, K. T.; Fredrickson, G. H.; Shea, J.-E.; Hwang, D. S.; Han, S. Dehydration Entropy Drives Liquid-Liquid Phase Separation by Molecular Crowding. Comm.Chem. 2020, 3 (1), 83. 10.1038/s42004-020-0328-8.

(54) André, A. A. M.; Yewdall, N. A.; Spruijt, E. Crowding-Induced Phase Separation and Gelling by Co-Condensation of PEG in NPM1-RRNA Condensates. Biophys. J. 2023, 122 (2), 397–407. 10.1016/j.bpj.2022.12.001.

(55) Strom, A. R.; Emelyanov, A. V; Mir, M.; Fyodorov, D. V; Darzacq, X.; Karpen, G. H. Phase Separation Drives Heterochromatin Domain Formation. Nature 2017, 547 (7662), 241–245. 10.1038/nature22989.

(56) Larson, A. G.; Elnatan, D.; Keenen, M. M.; Trnka, M. J.; Johnston, J. B.; Burlingame, A. L.; Agard, D. A.; Redding, S.; Narlikar, G. J. Liquid Droplet Formation by HP1α Suggests a Role for Phase Separation in Heterochromatin. Nature 2017, 547 (7662), 236–240. 10.1038/nature22822.

(57) Shakya, A.; King, J. T. DNA Local-Flexibility-Dependent Assembly of Phase-Separated Liquid Droplets. Biophys.J. 2018, 115 (10), 1840–1847. 10.1016/j.bpj.2018.09.022.

(58) Turner, A. L.; Watson, M.; Wilkins, O. G.; Cato, L.; Travers, A.; Thomas, J. O.; Stott, K. Highly Disordered Histone H1-DNA Model Complexes and Their Condensates. Proc. Natl. Acad. Sci. U. S. A. 2018, 115 (47), 11964–11969. 10.1073/pnas.1805943115.

(59) Vieregg, J. R.; Lueckheide, M.; Marciel, A. B.; Leon, L.; Bologna, A. J.; Rivera, J. R.; Tirrell, M. V. Oligonucleotide-Peptide Complexes: Phase Control by Hybridization. J.Am.Chem.Soc. 2018, 140 (5), 1632–1638. 10.1021/jacs.7b03567.

(60) Wang, L.; Gao, Y.; Zheng, X.; Liu, C.; Dong, S.; Li, R.; Zhang, G.; Wei, Y.; Qu, H.; Li, Y.; Allis, C. D.; Li, G.; Li, H.; Li, P. Histone Modifications Regulate Chromatin Compartmentalization by Contributing to a Phase Separation Mechanism. Mol.Cell 2019, 76 (4), 646–659.e6. 10.1016/j.molcel.2019.08.019.

(61) Dutagaci, B.; Nawrocki, G.; Goodluck, J.; Ashkarran, A. A.; Hoogstraten, C. G.; Lapidus, L. J.; Feig, M. Charge-Driven Condensation of RNA and Proteins Suggests Broad Role of Phase Separation in Cytoplasmic Environments. Elife 2021, 10. 10.7554/eLife.64004.

(62) Mimura, M.; Tomita, S.; Shinkai, Y.; Hosokai, T.; Kumeta, H.; Saio, T.; Shiraki, K.; Kurita, R. Quadruplex Folding Promotes the Condensation of Linker Histones and DNAs via Liquid-Liquid Phase Separation. J.Am.Chem.Soc. 2021, 143 (26), 9849–9857. 10.1021/jacs.1c03447.

(63) Soranno, A.; Incicco, J. J.; De Bona, P.; Tomko, E. J.; Galburt, E. A.; Holehouse, A. S.; Galletto, R. Shelterin Components Modulate Nucleic Acids Condensation and Phase Separation in the Context of Telomeric DNA. J.Mol.Biol. 2022, 434 (16), 167685. 10.1016/j.jmb.2022.167685.

(64) Leicher, R.; Osunsade, A.; Chua, G. N. L.; Faulkner, S. C.; Latham, A. P.; Watters, J. W.; Nguyen, T.; Beckwitt, E. C.; Christodoulou-Rubalcava, S.; Young, P. G.; Zhang, B.; David, Y.; Liu, S. Single-Stranded Nucleic Acid Binding and Coacervation by Linker Histone H1. Nat.Struct.Mol.Biol. 2022, 29 (5), 463–471. 10.1038/s41594-022-00760-4.

(65) Jack, A.; Kim, Y.; Strom, A. R.; Lee, D. S. W.; Williams, B.; Schaub, J. M.; Kellogg, E. H.; Finkelstein, I. J.; Ferro, L. S.; Yildiz, A.; Brangwynne, C. P. Compartmentalization of Telomeres through DNA-Scaffolded Phase Separation. Dev.Cell 2022, 57 (2), 277–290.e9. 10.1016/j.devcel.2021.12.017.

(66) Phan, T. M.; Kim, Y. C.; Debelouchina, G. T.; Mittal, J. Interplay between Charge Distribution and DNA in Shaping HP1 Paralog Phase Separation and Localization. Elife 2024, 12. 10.7554/eLife.90820.

(67) Nordenskiöld, L.; Shi, X.; Korolev, N.; Zhao, L.; Zhai, Z.; Lindman, B. Liquid-Liquid Phase Separation (LLPS) in DNA and Chromatin Systems from the Perspective of Colloid Physical Chemistry. Adv.Coll.Int.Sci. 2024, 326, 103133. 10.1016/j.cis.2024.103133.

(68) Dhakal, S.; Mondal, M.; Mirzazadeh, A.; Banerjee, S.; Ghosh, A.; Rangachari, V. α-Synuclein Emulsifies TDP-43 Prion-like Domain-RNA Liquid Droplets to Promote Heterotypic Amyloid Fibrils. Comm.Biol. 2023, 6 (1), 1227. 10.1038/s42003-023-05608-1.

(69) Dilweg, I. W.; Dame, R. T. Post-Translational Modification of Nucleoid-Associated Proteins: An Extra Layer of Functional Modulation in Bacteria? Biochem. Soc. Trans. 2018, 46 (5), 1381–1392. 10.1042/bst20180488.

