## Supporting Information for "Oligomerization-Mediated Phase-Separation in the Nucleoid-Associated Sensory Protein H-NS is Controlled by Ambient Cues"

AUTHOR INFORMATION

**Corresponding Authors**

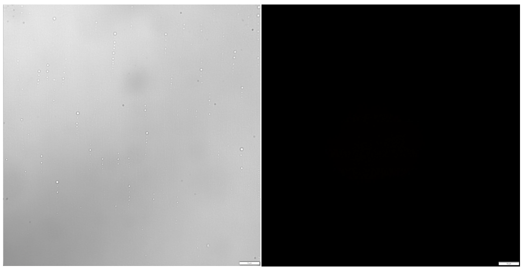

**Figure S1** DIC (left) and fluorescence microscopy (right) images of NHS-rhodamine-labeled 100 $\mu$M Y61E H-NS with 400 nM DNA. The scale bar (bottom right) represents 10 microns.

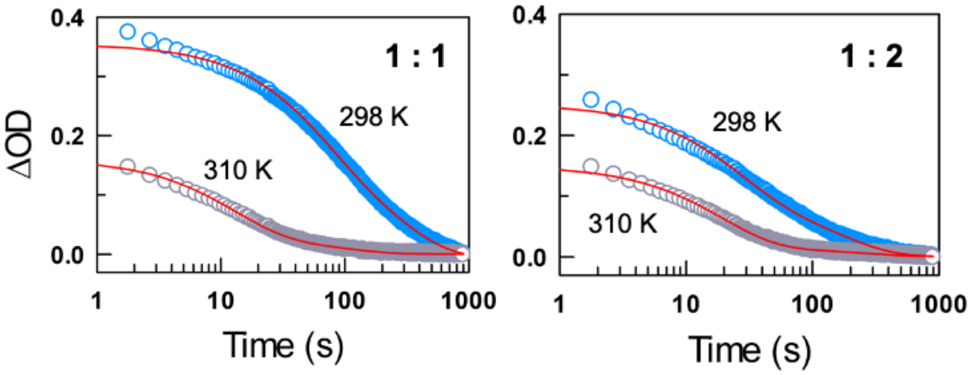


**Figure S2** Changes in optical density (OD) as a function of time upon mixing pre-formed H-NS:DNA condensates with Y61E at a molar ratio of 1:1 (left panel) or 1:2 (right panel) in a stopped-flow setup. Blue circles are experimental points while red curves are fits to bi-exponential functions. Note the poor fit at lower timescales necessitating the need for a tri-exponential function (see main text for details).

**Table S1** Parameters from tri-exponential fits to the data shown in main text Figure 5D, 5E.

| H-NS:Y61E Molar Ratio | Temperature | $\tau_1$ (s) | $a_1$ | $\tau_2$ (s) | $a_2$ | $\tau_3$ (s) | $a_3$ |
|---|---|---|---|---|---|---|---|
| 1:1 | 298 K | **2.3** ± 0.3 | 0.17 ± 0.02 | **50.5** ± 2.3 | 0.31 ± 0.01 | **210.0** ± 4.1 | 0.52 ± 0.01 |
| | 310 K | **1.5** ± 0.2 | 0.31 ± 0.02 | **16.2** ± 0.4 | 0.58 ± 0.02 | **114.1** ± 5.5 | 0.11 ± 0.01 |
| 1:2 | 298 K | **2.9** ± 0.2 | 0.29 ± 0.01 | **36.0** ± 0.8 | 0.45 ± 0.01 | **191.2** ± 3.8 | 0.26 ± 0.01 |
| | 310 K | **2.1** ± 0.3 | 0.27 ± 0.02 | **21.2** ± 0.5 | 0.64 ± 0.02 | **208.9** ± 14.4 | 0.09 ± 0.01 |
